# Resting-state functional connectivity and fast spindle temporal organization contribute to episodic memory consolidation in healthy aging

**DOI:** 10.1101/2024.11.28.625682

**Authors:** Anaïs Hamel, Pierre Champetier, Stéphane Rehel, Claire André, Brigitte Landeau, Florence Mézenge, Sacha Haudry, Daniel Roquet, Denis Vivien, Vincent de La Sayette, Gaël Chételat, Géraldine Rauchs, Alison Mary, the Medit-Ageing Research group

## Abstract

Episodic memory consolidation relies on the functional specialization of brain networks and sleep quality, both of which are affected by aging. Functional connectivity during wakefulness is crucial to support the integration of newly acquired information into memory networks. Additionally, the temporal dynamics of sleep spindles facilitates overnight memory consolidation by promoting hippocampal replay and integration of memories within neocortical structures. This study aimed at exploring how resting-state functional connectivity during wakefulness contributes to sleep-dependent memory consolidation in aging, and whether spindles clustered in trains modulates this relationship.

Forty-two healthy older adults (68.82 ± 3.03 years), enrolled in the Age-Well clinical trial, were included. Sleep-dependent memory consolidation was assessed using a visuo-spatial memory task performed before and after a polysomnography night. Resting-state functional connectivity data were analyzed using graph theory applied to the whole brain, specific brain networks and the hippocampus.

Lower limbic network integration and higher centrality of the anterior hippocampus were associated with better memory consolidation. Spindle trains modulated these effects, such that older participants with longer spindle trains exhibited a stronger negative association between limbic network integration and memory consolidation.

These results indicate that lower functional specialization at rest is associated with weaker memory consolidation during sleep. This aligns with the dedifferentiation hypothesis, which posits that aging is associated with reduced brain specificity, leading to less efficient cognitive functioning. These findings reveal a novel mechanism linking daytime brain network organization and sleep-dependent memory consolidation, and suggest that targeting spindle dynamics could help preserve cognitive functioning in aging.

## Introduction

One of the most common cognitive complaints among older adults is the growing difficulty in learning and remembering new information (Friedman et al., 2007; Tromp et al., 2015). Memory consolidation is a process that enables the stabilization of learned information and its transfer from the hippocampus to neocortical structures for long-term storage (Buzsáki, 1996). This process involves the repeated reactivation of memory traces and relies notably on the organization of brain networks and specific NREM sleep oscillations (i.e., slow oscillations and fast spindles) (Klinzing et al., 2019), both of which are known to undergo substantial age-related changes (Hamel et al., 2023; Muehlroth et al., 2020a; Nyberg, 2017). These alterations may contribute to the decline in episodic memory consolidation reported in numerous studies (see Muehlroth et al., 2020a for a review, but see also Sonni & Spencer, 2015; Wilson et al., 2012 for discrepant results).

The functional organization of brain networks has been proposed as a potential neural marker of cognitive plasticity (Gallen & D’Esposito, 2019), which may reflect the effectiveness of memory processes. Intrinsic or resting-state functional connectivity (rsFC) has a modular organization, characterized by specialized or segregated resting-state networks. Some studies revealed that these networks support distinct cognitive functions and that their topological organization varies with age (Damoiseaux, 2017; Deery et al., 2023). Efficient brain communication and cognition are supported by a fine balance between maintaining specialized brain networks (i.e., segregation) and facilitating interactions across distinct brain networks (i.e., integration) (Wig, 2017). Segregation minimizes wiring cost and supports flexibility and robustness in response to perturbations, while integration maximizes the efficiency of large-scale brain communication (Sporns & Betzel, 2016). Associative networks involved in high-level cognitive functions, such as the default mode (DMN), the frontoparietal control (FPCN) and the attentional networks are particularly vulnerable to aging and undergo a concomitant decrease in within-network connectivity (Betzel et al., 2014; Cassady et al., 2021; Chan et al., 2014; Damoiseaux, 2017; Geerligs et al., 2015) and an increase in between-network connectivity (Betzel et al., 2014; Cassady et al., 2021; Ferreira et al., 2016; Grady et al., 2016; Malagurski et al., 2020; Spreng et al., 2016) with advancing age. This is consistent with the dedifferentiation hypothesis of cognitive aging, which assumes that decreased functional specialization leads to more diffuse brain recruitment, less distinct neural representations and reduced processing efficiency, which may contribute to cognitive decline (Baltes & Lindenberger, 1997; Koen & Rugg, 2019; Reuter-Lorenz & Lustig, 2005).

In older adults, better episodic memory performance has been associated with higher rsFC within the DMN and the FPCN (Huo et al., 2018; Shaw et al., 2015). Besides within-network changes, age-related increase in FC between the FPCN, the dorsal attentional network and the DMN may impair flexibility in allocating attentional resources (Esposito et al., 2018; Grady et al., 2016; Spreng et al., 2016). However, the association between cognition and the age-related increase in between-network FC remains unclear. Some studies found that this reduced network specialization is associated with lower cognitive performance (Chan et al., 2014; Huo et al., 2018; Pedersen et al., 2021; Shaw et al., 2015; Stumme et al., 2020), while others have reported an association with better cognition (Crowell et al., 2020; Grady et al., 2016). To explain these discrepancies, it was proposed that stronger between-network FC associated with better cognition might reflect a compensatory process, whereas the same association with lower between-network FC might indicate less dedifferentiation (Deery et al., 2023). Despite the inconsistent evidence regarding the association between network integration and cognition, studies that specifically examined memory consolidation provided valuable perspectives. Two studies have examined wake-dependent memory consolidation by investigating post-learning changes in rsFC in young and older adults (Jacobs et al., 2015; Kukolja et al., 2016). Efficient memory consolidation was associated with post-encoding rsFC changes in brain regions specific to learning (Kukolja et al., 2016). Compared to younger adults, older adults showed a reduced ability to increase connectivity in the right lingual gyrus as part of an extended DMN during this post-encoding stage (Kukolja et al., 2016). The other study, using the same dataset, reported that post-encoding memory consolidation in older adults was more strongly associated with between-networks rsFC, suggesting that in older adults, memory consolidation rely more on between-network connectivity than within-network or region-specific connectivity (Jacobs et al., 2015). Despite growing evidence that rsFC may help explain age-related differences in memory performance (Fjell et al., 2015, 2016), no studies to date have explored how rsFC relates to sleep-dependent episodic memory consolidation in older adults. Moreover, it is essential to use methodologies that provide a comprehensive view of brain organization to further our understanding of the relationship between rsFC and memory consolidation.

Graph theory is an increasingly relevant approach in the field of aging to identify associations between brain network features and cognition. Graph theory conceptualizes the brain as a complex network, where nodes represent brain regions of interest (ROIs) and edges represent the connections between them (Bullmore & Sporns, 2009; Rubinov & Sporns, 2010). This complex network model provides a complete overview of the rsFC patterns at multiple levels, including whole-brain, networks and ROIs. Studies using graph theory showed age-related disruptions in the modular/segregated brain organization (Betzel et al., 2014; Cao et al., 2014; Damoiseaux, 2017; Geerligs et al., 2015, 2015; Onoda & Yamaguchi, 2013) leading to decreased global efficiency, whole-brain modularity and segregation along with increased integration of associative networks (Chan et al., 2014; Geerligs et al., 2015; Sala-Llonch, 2014; Song et al., 2014). The application of graph theory represents a promising method for understanding the potential associations between functional networks and sleep-dependent memory consolidation.

Besides functional connectivity, sleep also plays a crucial role in cognitive functioning, particularly memory consolidation. Sleep is an optimal physiological state for stabilizing recent and labile memory traces and strengthening synaptic connections (Diekelmann & Born, 2010). For episodic memories, this process is notably supported by non-rapid eye movement (NREM) sleep oscillations, which initiate the reactivation of hippocampal traces and their subsequent integration into neocortical structures, a mechanism known as the hippocampo-neocortical dialogue (Buzsáki, 1996). Fast sleep spindles (13-15Hz) were proposed to promote the reactivation of hippocampal representations and synaptic plasticity (Cairney et al., 2018) through their temporal coordination with slow oscillations and sharp-wave ripples (Born & Wilhelm, 2012; Hamel et al., 2023 for a review). During aging, sleep changes play a significant role in memory consolidation decline (Muehlroth et al., 2020a). For instance, sleep spindles show decreases in density, duration, and amplitude (Crowley, 2002; Fernandez & Lüthi, 2020; Landolt et al., 1996; Nicolas et al., 2001). Interestingly, while fast spindles are generally believed to support memory consolidation in young adults (Cairney et al., 2018), evidence of such a role in older individuals is less convincing. For instance, studies conducted in older adults have not consistently reported an association between fast spindle density and memory consolidation using either a scene-word task (Muehlroth, et al., 2020b) or an object-location association task (Champetier et al., 2023). These results suggest that the relationship between spindles and memory consolidation may be more complex in aging, potentially involving other factors than spindle density. Recently, it has been shown that sleep spindles exhibit specific temporal dynamics, occurring either in isolation or very close in time, forming clusters called “spindle trains” (Boutin & Doyon, 2020). In a previous study from our group, Champetier et al. (2023) investigated the age-related modifications of fast spindle trains as well as its impact on memory consolidation in older adults. We showed that the mean size of fast spindle trains decreased after 50 years and was positively associated with memory consolidation. This finding supports the idea that the repetitive occurrence of spindles may facilitate the reactivation and reorganization of newly encoded memories within brain networks during sleep (Boutin & Doyon, 2020). However, while the role of spindle dynamics during the overnight reorganization of memory traces is becoming increasingly understood, the association with functional brain organization remains unclear. Given their role in memory consolidation, sleep spindles might also influence the association between rsFC and memory consolidation.

The present study assesses, for the first time, the association between rsFC and sleep-dependent memory consolidation, in cognitively unimpaired older adults. We used complementary graph metrics to measure segregation and integration at both the whole-brain and network levels as well as centrality at the hippocampal level. According to the dedifferentiation hypothesis, we surmised that better sleep-dependent memory consolidation would be associated with higher segregation and lower integration of associative networks, as well as higher hippocampal centrality, reflecting the key role of this structure in the hippocampo-neocortical dialogue. We also examined the rsFC of the anterior and posterior hippocampus, given their respective contributions to the processing of object/item and spatial/contextual information (Ranganath & Ritchey, 2012). Finally, given that higher number of fast spindles per train has been associated with better overnight memory consolidation in aging (Champetier et al., 2023), we also aimed at evaluating the potential modulatory role of spindle trains in the relationships between functional connectivity and memory consolidation.

## Materials and methods

### Participants

A total of 42 cognitively unimpaired older adults were included in this study, corresponding to part of the baseline data of the Age-Well randomized controlled clinical trial (RCT) of the Medit-Ageing European Project (Poisnel et al., 2018) sponsored by the French National Institute of Health and Medical Research (INSERM) (**Figure 1**). Participants’ demographical variables, sleep parameters and rsFC measures are provided in **Table 1** and supplementary **Table S1**. The study included community-dwelling individuals aged over 65 years, with normal cognitive performance based on standardized cognitive tests. Participants were native French speakers, retired for at least one year, with a minimum of 7 years of education. Exclusion criteria encompassed safety concerns with MRI or PET, major neurological or psychiatric disorders (including substance abuse), history of cerebrovascular disease, chronic illness, acute instability, and medication affecting cognitive functions. Participants underwent an assessment that included a comprehensive neuropsychological evaluation, in-home polysomnography (PSG), structural and functional MRI, all these data being collected within a maximum period of 3 months (**Figure 2A**). Participants provided their written informed consent prior to the examinations. The Age-Well RCT received approval from the ethics committee (CPP Nord-Ouest III, Caen; trial registration number: EudraCT: 2016-002441-36; IDRCB: 2016-A01767-44; ClinicalTrials.gov Identifier: NT02977819).

**Figure 1.**
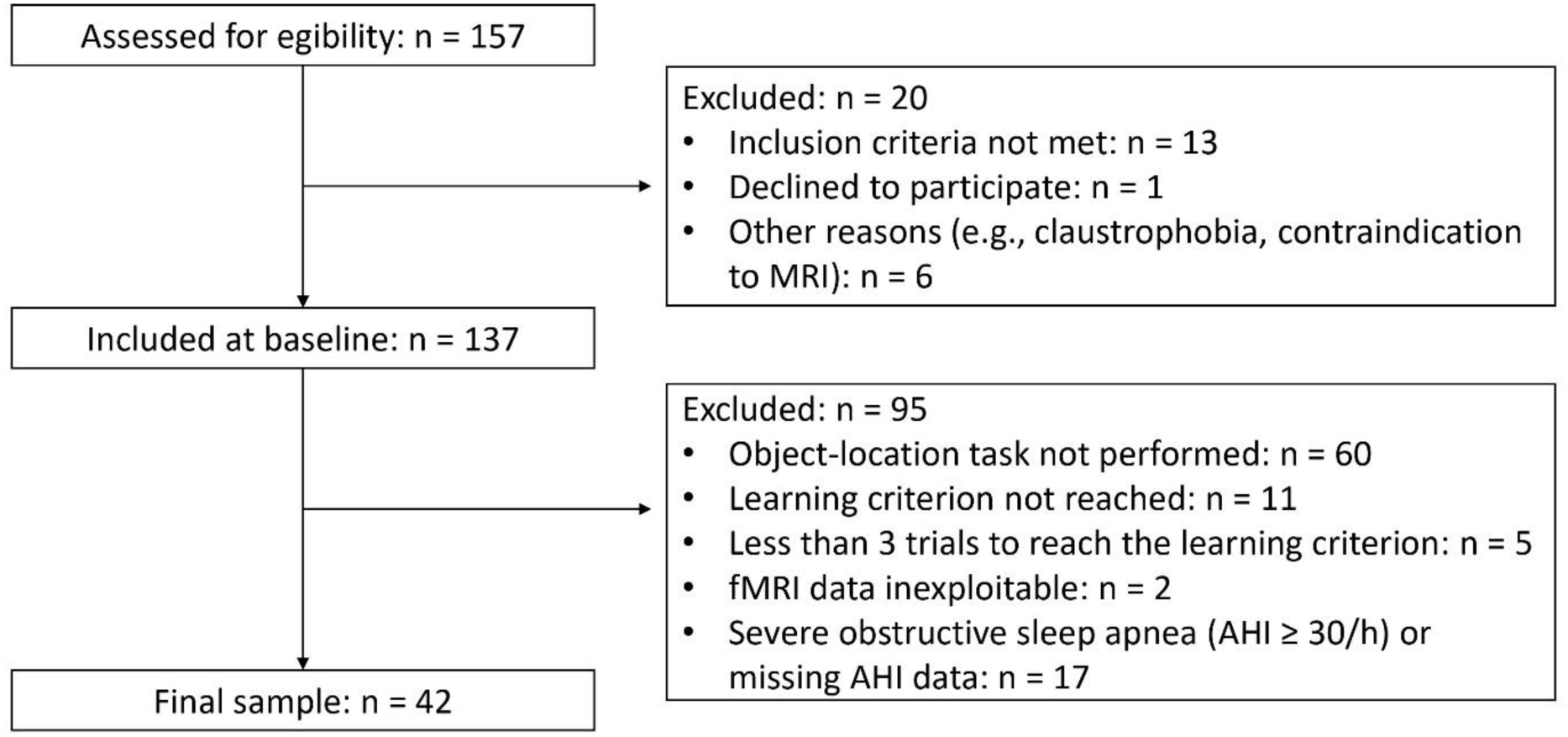
Flow diagram of the inclusion process.

**Table 1.** Participants’ characteristics (n = 42).

| Demographic |  |  |
| --- | --- | --- |
| Age (years) |  | 68.82 ± 3.03 [65:77] |
| Sex ratio (F/M) |  | 28/14 |
| Education (years) |  | 12.98 ± 3.14 [7:20] |
| Mini-Mental State Examination (/30) |  | 29.26 ± 0.83 [27:30] |
| Sleep |  |  |
| Total sleep time (TST; min) |  | 361.13 ± 67.66 [227:493.5] |
| Sleep onset latency (min) |  | 21.85 ± 15.31 [2:64.5] |
| Wake after sleep onset (min) |  | 86.74 ± 47.38 [13.5:230] |
| Sleep efficiency (%) |  | 76.90 ± 16.44 [52.5:91] |
| NREM-1 sleep (% of TST) |  | 11.02 ± 4.00 [4.7:21.9] |
| NREM-2 sleep (% of TST) |  | 48.69 ± 7.58 [29.7:63.5] |
| NREM-3 sleep (% of TST) |  | 20.14 ± 8.65 [1.9:48.1] |
| REM sleep (% of TST) |  | 20.15 ± 5.12 [5.1:33.3] |
| Apnea-Hypopnea Index (number of events/hour) |  | 18.72 ± 6.92 [4:29.9] |
| Mean number of fast spindles per train |  | 3.08 ± 0.31 [2.60:3.85] |

Continuous variables are indicated as follows: mean ± SD [min, max]; *TST*: Total Sleep Time.

**Figure 2.**
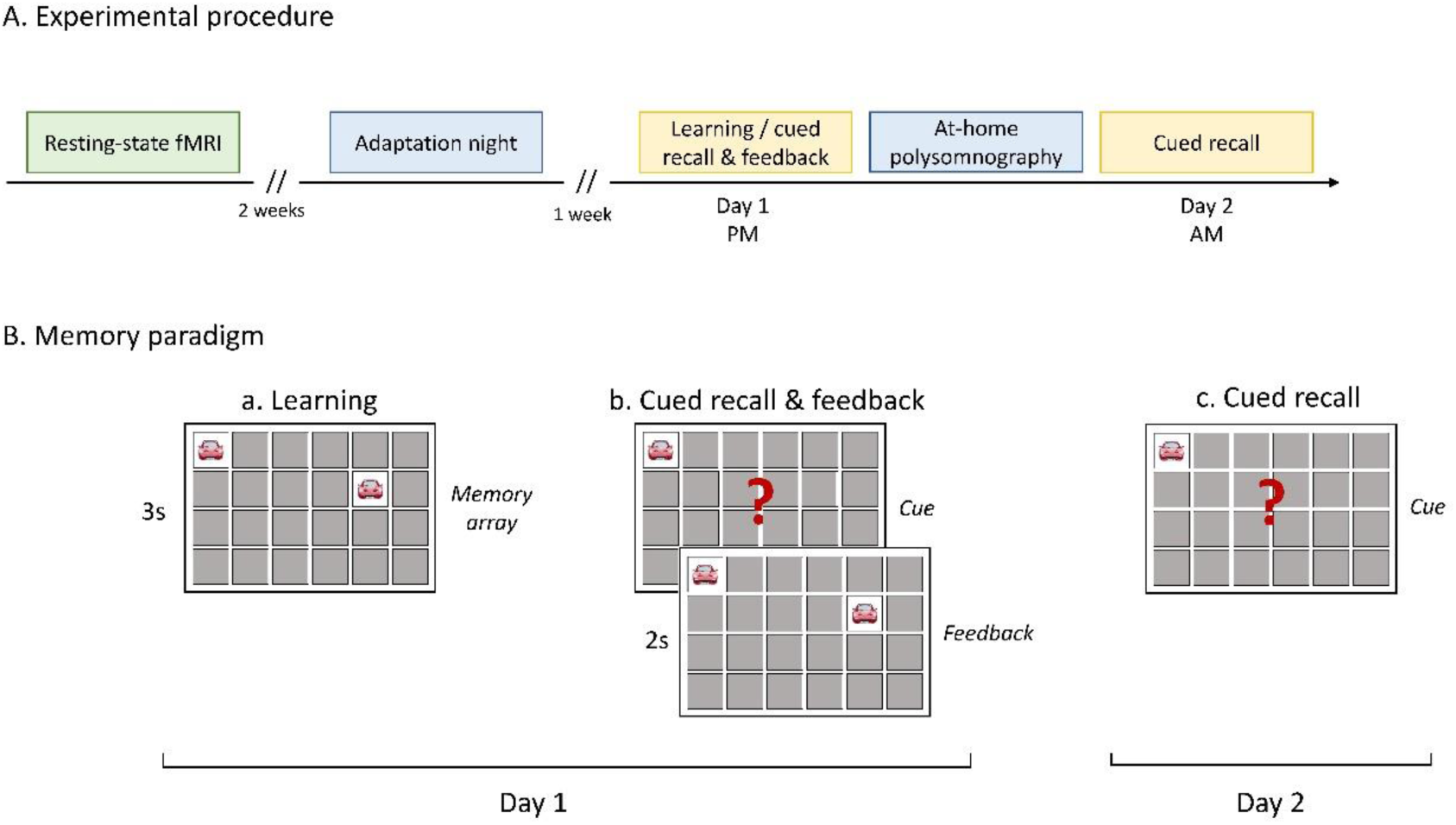
Study design and memory task. **(A) Experimental procedure.** Resting-state functional data were collected on average 2 weeks prior to the habituation night (resting-state functional data acquisition – habituation night [range: -23 to 62 days]). Participants underwent an adaptation night to familiarize them with the sleep recording equipment approximately one week before the experimental night. The memory task consisted of a learning phase and cued recall with feedback on Day 1 and a cued recall on Day 2. Sleep was monitored using in-home polysomnography on post-learning night**. (B) Object-location memory task**: (a) During the learning phase, participants were exposed to each pair of cards as follows: the first card of each pair was presented for one second, then both cards were displayed simultaneously for three seconds. The 12 card-pairs were presented twice in the same order, with a three-second inter-stimulus interval between each pair. These two runs were followed by a cued recall procedure (b) in which the first card of each pair was presented and participants were asked to use a computer mouse to indicate the location of the second card. Regardless of whether the answer was correct or not, a visual feedback was provided by displaying the second card in the correct location for two seconds. The cued recall procedure continued until participants reached a criterion of 66.7% correct responses in a trial (i.e. 8 card pairs correctly located). The last score obtained was defined as the learning score. This learning phase was performed in the late afternoon before the polysomnography night. (c) Memory recall was assessed after a night of sleep following the same procedure as on Day 1, but with a single trial without feedback.

### Visuo-spatial memory task

Participants performed a computerized visuospatial object-location task adapted from Rasch et al. (2007) and known to be hippocampus-dependent and reinforced by sleep (Rasch et al., 2007). This task was fully described in Champetier et al. (2023). Briefly, participants were invited to learn, in the late afternoon, the locations of 12 card pairs on a computer screen until reaching a criterion of 66.7% correct responses (Figure 2B). At retrieval testing after a night of sleep recorded with in-home PSG, memory of the card locations was assessed using a single trial.

Overnight memory consolidation was calculated as follows: [(recall score – learning score) / learning score] x 100. Of note, 11 participants did not reach the learning criterion and were excluded from the analyses. Sixty participants did not perform the task due to a late implementation in the protocol or time constraints. To ensure a minimum level of practice and repetition during the learning phase, participants who reached the learning criterion in less than 3 trials were also removed (n = 5).

### Polysomnography recording

PSG was performed in-home using a portable recording device (Siesta®, Compumedics, Australia). All the participants included in the analyses underwent a habituation night one week before the experimental night. The PSG montage included an electroencephalogram (EEG), electrooculogram (EOG), electrocardiogram (ECG), and chin electromyogram (EMG). Respiratory movements, airflow and oxygen saturation were also recorded to score respiratory events. For the EEG recording, 20 electrodes were placed over the scalp in line with the international 10-20 system (Fp1, Fp2, F3, F4, F7, F8, FZ, C3, C4, CZ, T3, T4, P3, P4, PZ, O1, O2, vertex ground and bi-mastoid reference), with impedances kept below 5 kΩ. The EEG signal was digitalized at 256 Hz, high-pass at 0.3 Hz and low-pass filters at 35 Hz were applied. Recordings were visually scored in 30-second epochs following the international scoring rules of the American Academy of Sleep Medicine. Sleep apnea was characterized as a reduction in airflow ≥ 90% for at least 10 seconds. Sleep hypopnea was defined as a reduction in airflow ≥ 30% lasting ≥ 10 seconds, associated with an arousal or a ≥ 3% oxygen desaturation. Eight participants had an apnea-hypopnea index (AHI, the sum of apneas and hypopneas per hour of sleep) ≥ 30 events/h, indicating severe obstructive sleep apnea (OSA), and one with missing AHI data were removed from the analyses.

The method used for fast spindle detection and train identification was described in Champetier et al., 2023. Briefly, fast spindles were automatically detected in artifact-free N2-N3 epochs using a subject-specific frequency band. First, for each subject, individual frequency peak in the fast spindle range was visually identified on centroparietal channels and considered as the fast spindle center frequency (mean ± SD = 13.66 ± 0.05 Hz). Then, the EEG signal was filtered with a band-pass width of 2 Hz centered on the fast spindle center frequency. A fast spindle was detected when the smoothed root mean square signal exceeded an individual threshold of 1.5 standard deviations of the filtered signal for 0.5-2 seconds. Fast spindles were excluded when amplitude exceeded 120 µV.

Spindle trains were calculated on each channel separately and consisted of at least two fast spindles interspaced by less than 6 seconds. The number of fast spindles per train, also referred as length of spindle trains, was calculated separately on C3 and C4 and then averaged for each participant.

### Neuroimaging acquisition

Neuroimaging data were collected on average 2 weeks prior to the habituation night (mean ± SD = 20 ± 14 days). Imaging data were collected using a Philips Achieva 3T Philips (Eindhoven, The Netherlands) scanner at the Cyceron Center (Caen, France). Different MRI sequences were used for the current study: high-resolution T1-weighted structural images were collected (3D-T1-FFE sagittal; repetition time = 7.1 ms; echo time = 3.3 ms; flip angle = 6°; 180 slices with no gap; slice thickness = 1 mm; field of view = 256×256 mm^2^; in plane resolution = 1×1 mm^2^). Then, resting-state functional volumes were obtained (2D-T2*-FFE-EPI axial, SENSE = 2.5; Time Repetition = 2400 ms; Time Echo = 30 ms; flip angle = 85°; 44 slices with no gap; slice thickness = 2.8 mm; field of view = 200×200 mm^2^; in-plane resolution = 2.5×2.5 mm^2^; 200 volumes). Participants were equipped with earplugs and their heads were stabilized with foam pads to minimize head motion. During the 8-min resting-state acquisition, participants were instructed to keep their eyes closed, let their mind wander and not fall asleep. Alertness during the scan was confirmed by post-scan debriefing.

### Pre-processing resting-state fMRI data

Data were pre-processed using SPM12 toolbox (Wellcome Trust Centre for Neuroimaging, https://www.fil.ion.ucl.ac.uk/spm/software/spm12/) implemented in MATLAB R2018b. Rs-fMRI data were checked for motion artefacts or for abnormal variance distribution using the TSDiffAna routine (https://imaging.mrc-cbu.cam.ac.uk/imaging/DataDiagnostics). One participant showing significant movement (> 3mm translating or 2° rotating) was excluded for further analysis. All EPI volumes were slice timing corrected, realigned to the first volume and a data denoising process was conducted. Initially, an individual independent component analysis (ICA) was performed on each participant’s functional MRI FSL’s MELODIC (Multivariate Exploratory Linear Decomposition into Independent Components). This process decomposed the fMRI data into 50 spatially independent components per participant. Each component was visually examined, and those showing stripes alternating along the z-axis were identified as noise. Subsequently, these noise components were removed from the data using fsl_regfilt command by regressing out their time courses from the fMRI data. All data were normalized within the native space to correct for distortion effects (Villain et al., 2010). Then, fMRI data were co-registered to the T1-weighted MRI images, normalized to the MNI 1.5 mm space by applying the normalization parameters derived from the segmented T1 images, and smoothed with a 4-mm full-width at half-maximum Gaussian kernel. Finally, a temporal band-pass filter between 0.01 and 0.08 Hz and a mask including only gray matter voxels were applied to the images.

### Parcellations selection

The cortical and hippocampal surface of each participant was parcellated using a combination of the Schaefer cortical (Schaefer et al., 2018) and the Melbourne subcortical (Tian et al., 2020) atlases, which are both based on rsFC data. These atlases provide a fine topographic organization of human cortical and subcortical areas, and have been used in studies investigating rsFC (Deleglise et al., 2023; Wu et al., 2022). The cortical structure was divided into 100 anatomical ROIs using the 100-parcel Schaefer atlas. Each ROI was assigned to one of the 7 putative networks including the dorsal attention, salience/ventral attention, limbic, frontoparietal control, default mode, visual, sensorimotor networks (Schaefer et al., 2018). Additionally, the “hippocampal network” included the left and right anterior/posterior hippocampal regions from the 32-parcel Melbourne Atlas. The combination of the Schaefer and Melbourne parcellations resulted in 104 ROIs assigned to one of 8 networks (Figure 3A).

**Figure 3.**
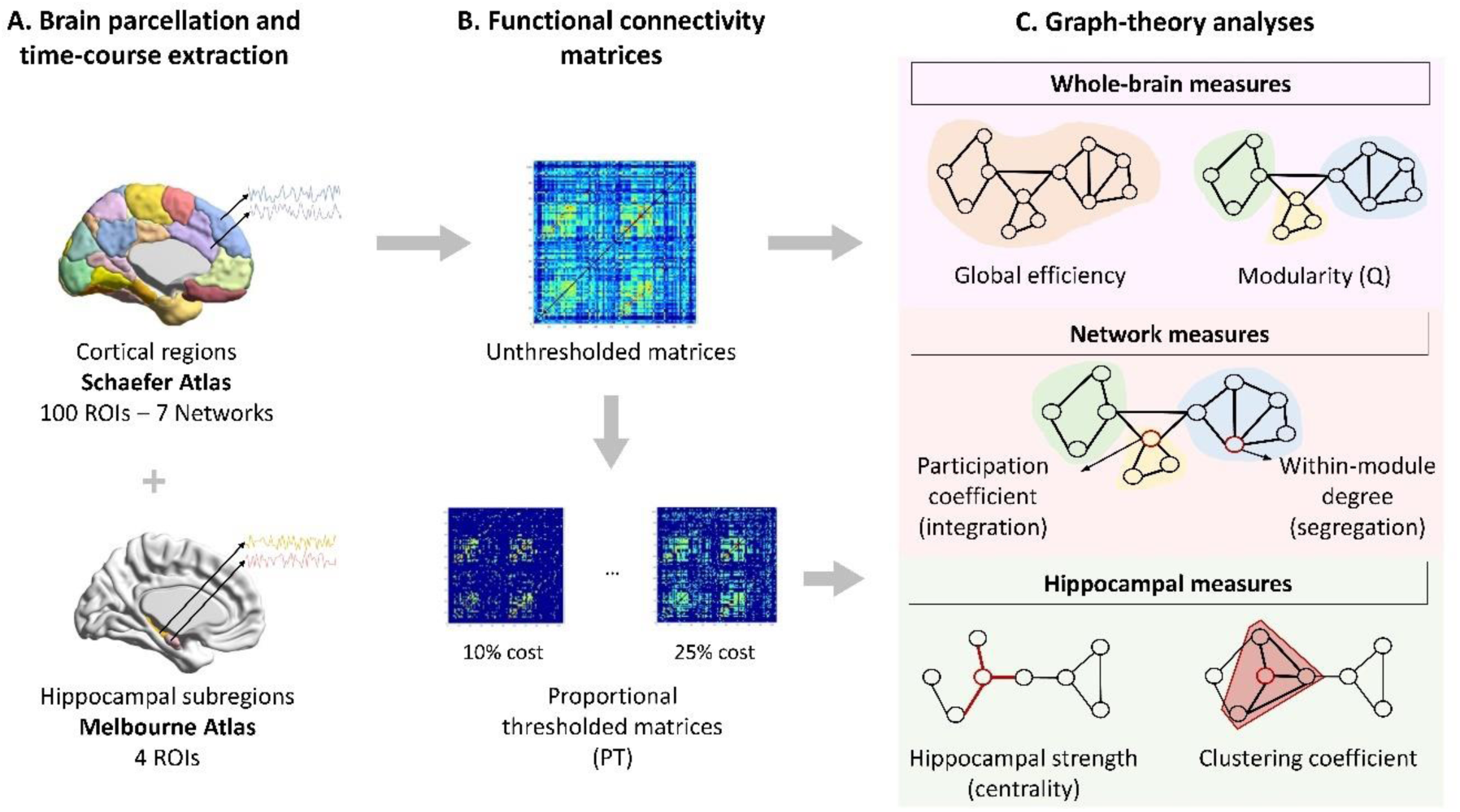
Overview of functional connectivity analyses applied to resting-state functional MRI data. (A) Brain parcellation of Schaefer atlas and hippocampal subregions from the Melbourne atlas. BOLD time-course were averaged within each ROI. (B) The Pearson’s correlation (Fisher’s r-to-z transformation) was used to assess functional connectivity between each pair of ROIs. A proportional threshold of 10% to 25% (incremented by 5) of the strongest connections was applied to the connectivity matrices. (C) Summary of graph measures. Whole-brain measures included global efficiency, which quantifies the efficiency of information transfer across the whole-brain, and modularity, which indicates the balance between functional connectivity within- and between-networks across the whole-brain. For network measures, the participation coefficient indicates the degree of between-network FC (integration) and the within-module degree quantifies the degree of within-network FC (segregation). Hippocampal measures included the hippocampal strength which quantifies the centrality of the hippocampus based on the strength of connectivity between the hippocampus and all other ROIs in the whole-brain. We also computed the clustering coefficient, which indicates the extent to which the interconnection of the hippocampus with its neighbors forms a cluster.

After a co-registration of the atlases to the 1.5 mm fMRI data space, Pearson’s correlations between each pair of ROIs time series were extracted using the Brainnetome fMRI Toolkit (https://sphinx-doc-brant.readthedocs.io/en/latest/index.html, Xu et al., 2018). Then, the 104 x 104 ROI-to-ROI connectivity matrices generated for each participant were converted to normalized z-scores using Fisher transformation.

### Functional connectivity matrices threshold

Following a Fisher’s z transformation of FC matrices, the diagonal and all negative weights of the connectivity matrix were adjusted to zero. Negative weights are usually removed in the matrix (Chan et al., 2014) due to the ambiguity of the meaning and interpretation of negative correlations A proportional threshold was applied to the connection matrices to include 10 to 25% of the strongest connections (incremented by 5; Figure 3B), as in previous studies (Cohen & D’Esposito, 2016; Luppi et al., 2019). Matrix thresholding is recommended when using graph theory approaches to improve robustness and avoid false positive connections (Zalesky et al., 2016).

### Graph analyses

Whole-brain, network and hippocampal FC were extracted using the Brain Connectivity Toolbox (see Supplementary methods for details, and Figure 3C). Whole-brain measures included global efficiency, which quantifies the efficiency of information transfer across the whole-brain, and modularity, which indicates the balance between functional connectivity within- and between-networks across the whole-brain. For network measures, the participation coefficient indicates the degree of between-network FC (integration) and the within-module degree quantifies the degree of within-network FC (segregation). Hippocampal measures included hippocampal strength which quantifies the centrality of the anterior/posterior hippocampal subregion based on the strength of connectivity between the anterior/posterior hippocampus and all other ROIs in the whole-brain and clustering coefficient, which indicates the extent to which the interconnection of the anterior/posterior hippocampal with its neighbors forms a cluster.

### Statistical analyses

Statistical analyses were conducted with the R statistical software package (version 4.1.1). The assumptions for linear regression were tested, with 99% meeting normality and 97.5% meeting homoscedasticity. Parametric tests were used as the sample size (n = 42) satisfied the central limit theorem. Associations between overnight memory consolidation and rsFC were examined using multiple linear regression including whole-brain measures, Schaefer’s Atlas network measures (7 networks), and anterior and posterior hippocampal ROIs measures. Memory consolidation was treated as the dependent variable, and graph measures as the independent variables. Age, sex, education and AHI were included as covariates of no interest. For significant associations found with network integration measures, network segregation (modularity (Q)) was included as a covariate in the statistical models, given the strong association between increasing integration and decreasing network segregation that has been observed in aging (Chan et al., 2014). To confirm these results, the analyses were also performed using an alternative measure of system segregation implemented by Chan et al., 2014. This measure was calculated as the difference between mean within-network FC and mean between-network FC divided by mean within-network FC (Chan et al., 2014). To investigate the statistical associations between graph measures and memory consolidation, a series of complementary analyses were conducted to verify that the variables of the statistical model were not associated with other graph measures and that graph measures were not associated with each other (see Supplementary methods and results and **Tables S2 and S3**). Then, for any significant association between memory consolidation and rsFC, we also examined the possible interaction effect of the mean number of fast spindls per train. These analyses were repeated for graph measures derived from the unthresholded and proportional thresholded matrices. Results were reported at the 15% threshold as it represented the lowest threshold at which all findings were consistent, ensuring a robust and coherent basis for our analysis while capturing higher specificity (Zalesky et al., 2016). All p-values were FDR-corrected to control for Type I errors due to multiple comparisons (Benjamini & Hochberg, 1995).

## Results

### 1. Association between rsFC and memory consolidation

#### Whole-brain analyses

No significant associations were found between memory consolidation and global efficiency or modularity (Q) (supplementary **Table S4**).

#### Network analyses

We found a significant association between memory consolidation and the participation coefficient of the limbic network (supplementary **Table S4**). Higher overnight memory consolidation was associated with lower median participation coefficient of the limbic network (β = -0.46, p = 0.049, FDR corrected, Figure 4).

**Figure 4.**
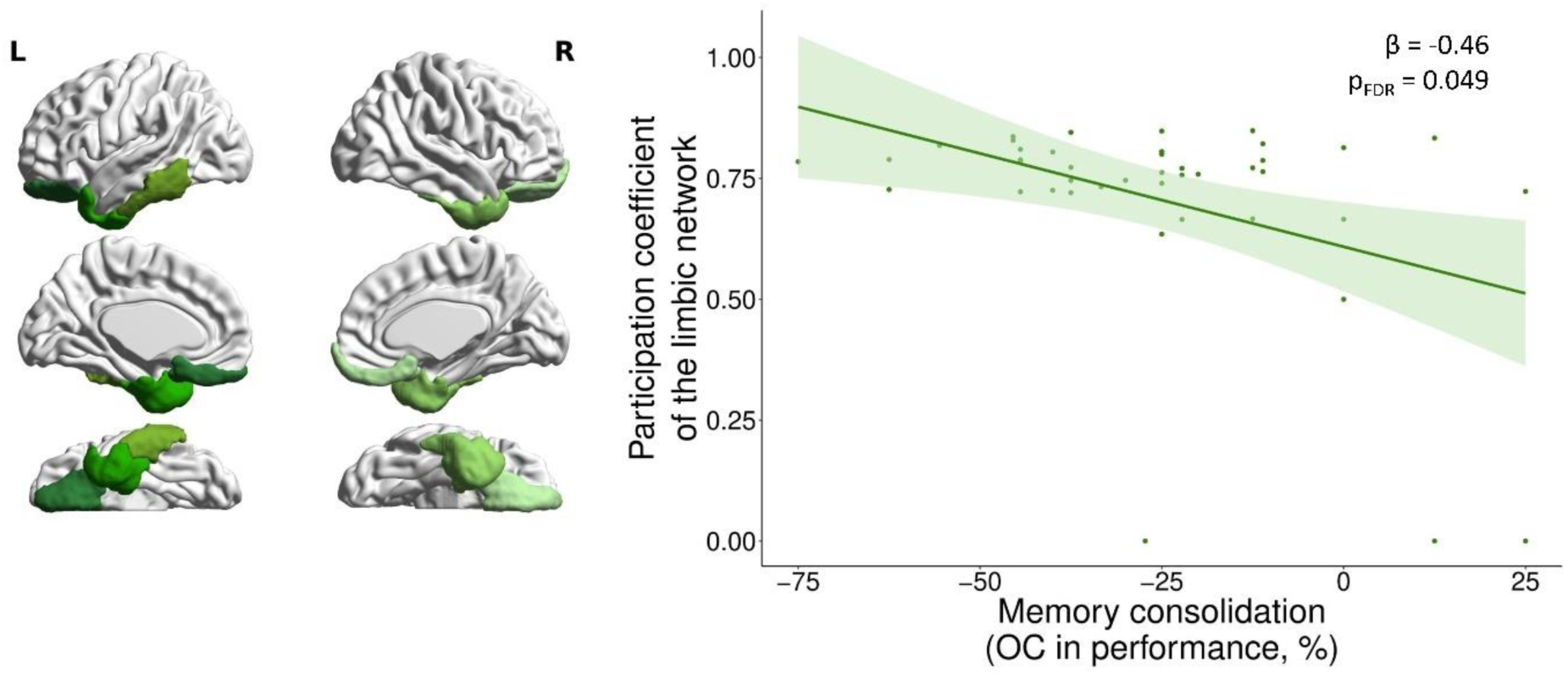
Association between overnight memory consolidation and between-network functional connectivity (participation coefficient) of the limbic network. Visual representation of the limbic network (left panel) and regression between memory consolidation and limbic network integration observed at p = 0.049, FDR corrected (right panel). Analyses were adjusted for age, sex, education and AHI. *FDR: False Discovery Rate; L: Left; OC: Overnight Change; R: Right*.

Complementary analyses were performed by adding a whole-brain segregation measure as covariate in the model. The negative association between memory consolidation and limbic network integration remained significant even when controlling by the modularity index (Q) (β = -0.44, p= 0.009) or system segregation measure (β = -0.43, p = 0.014) (see Methods and supplementary **Table S5**)

No significant association was found between memory consolidation and within-module degree of networks (supplementary **Table S4**).

#### Hippocampal analyses

Memory consolidation was positively associated with the functional centrality (strength) of the anterior (β = 0.40, p = 0.046, FDR corrected) but not of the posterior hippocampus (Figure 5). No significant association was found between the clustering coefficient of the anterior or posterior hippocampus and memory consolidation (supplementary **Table S4**).

**Figure 5.**
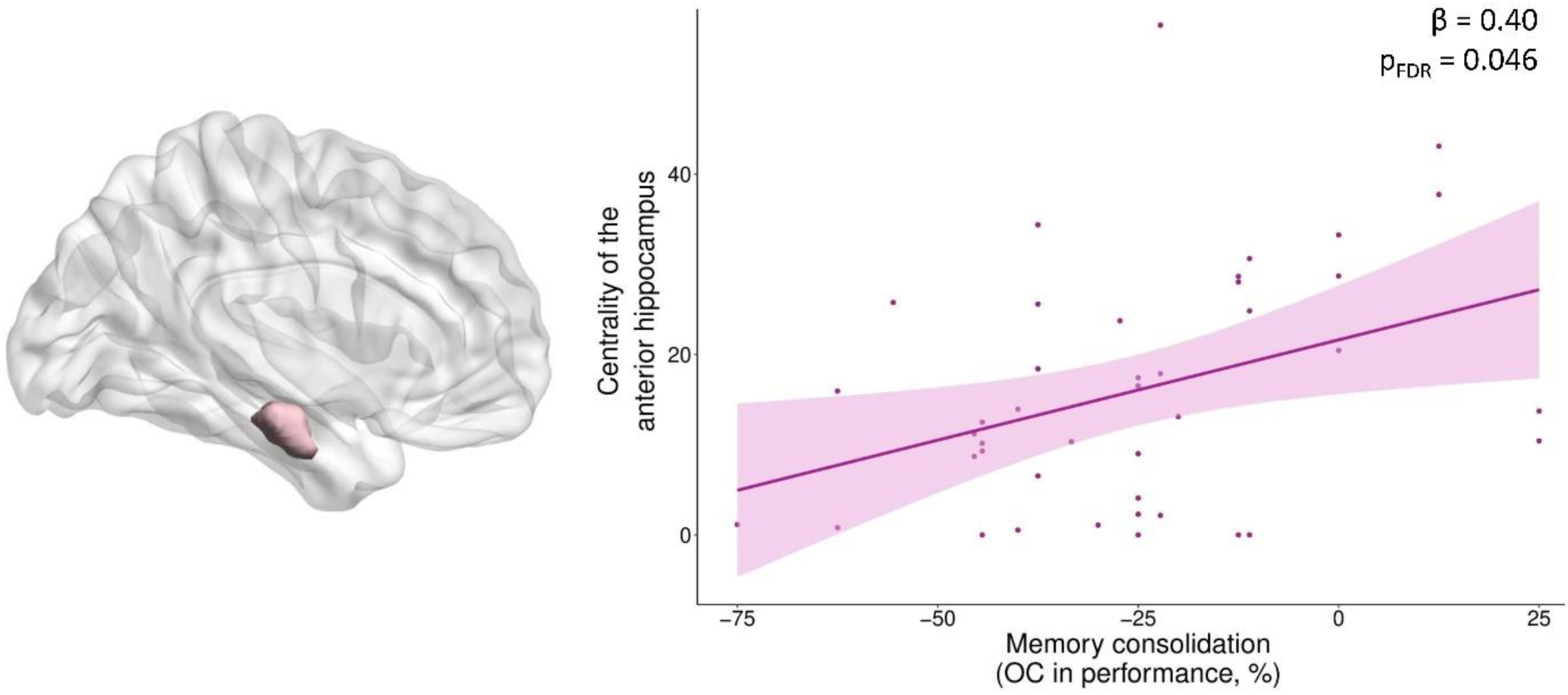
Association between overnight memory consolidation and the functional centrality (strength) of the anterior hippocampus. Visual representation of the anterior hippocampus (left panel) and regression between memory consolidation and anterior hippocampus centrality observed at p = 0.046, FDR corrected (right panel). Analyses were adjusted for age, sex, education and AHI. *FDR: False Discovery Rate; OC: Overnight Change*.

### 2. Association between rsFC and memory consolidation: modulation by the mean number of fast spindles per train

The mean number of fast spindles per train modulated the association between memory consolidation and the participation coefficient of the limbic network. Specifically, the association between the limbic network integration and overnight memory consolidation was found for participants with longer spindle trains (β = -3.45, p = 0.02; Figure 6). For graphical representation of interactions, participants were grouped according to whether their mean number of fast spindles per train was above or below the median value of 3.03 fast spindles per train. Mean number of fast spindles per train did not modulate the association between overnight memory consolidation and the functional centrality of the anterior hippocampus (supplementary **Table S6**).

**Figure 6.**
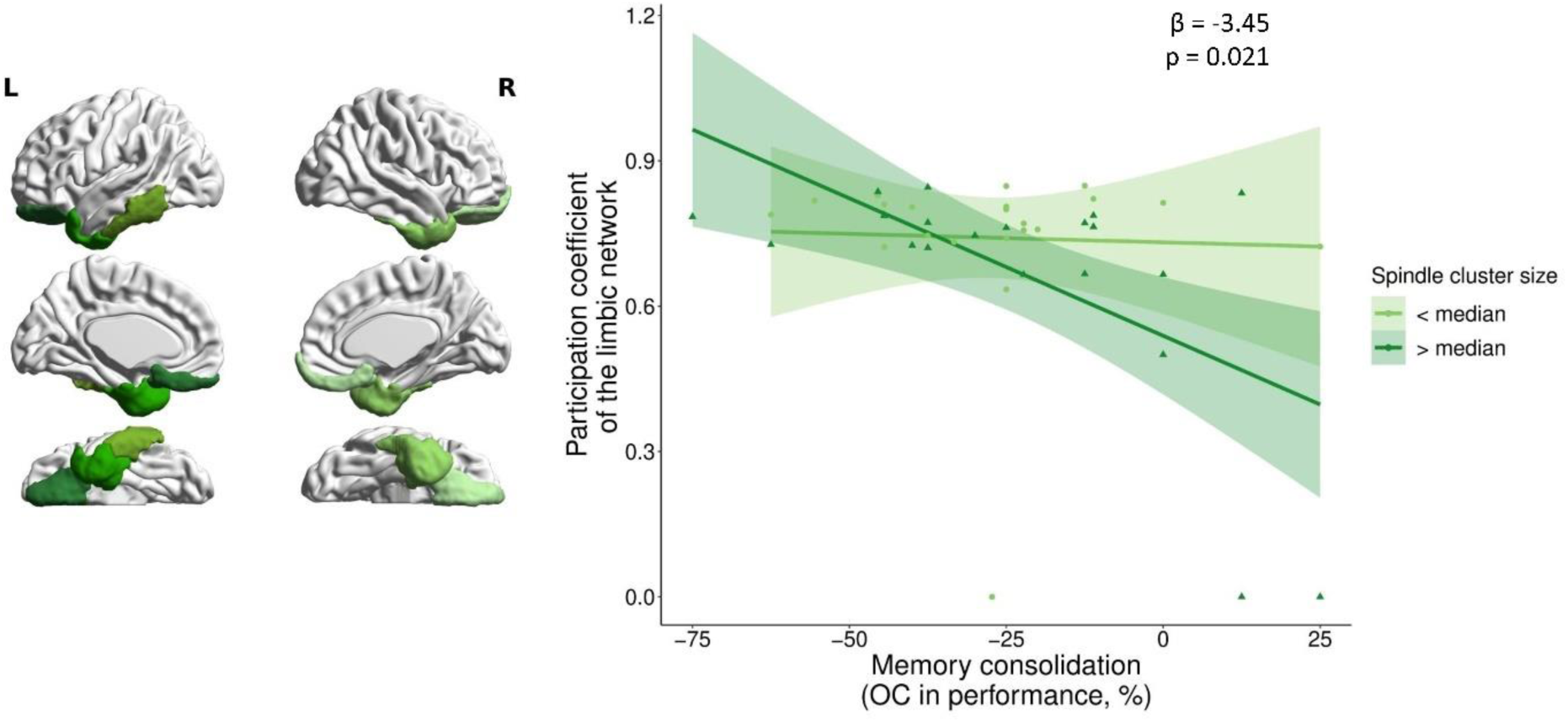
Modulation of the association between memory consolidation and between-network FC (participation coefficient) of the limbic network by the mean length of spindle trains. Visual representation of the limbic network (left panel), a modulation effect of fast spindle trains on associations between resting-state functional connectivity and memory consolidation was observed at p = 0.02 (right panel). Analyses were adjusted for age, sex, education and AHI. *FDR: False Discovery Rate; L: Left; OC: Overnight Change; R: Right*.

## Discussion

The present study investigated the associations between sleep-dependent memory consolidation and intrinsic functional organization in older adults, highlighting the critical role of network integration (participation coefficient) and hippocampal centrality. Our analyses revealed that lower between-network FC of the limbic network and higher anterior hippocampal centrality were associated with better memory consolidation. Additionally, the temporal organization of fast sleep spindles modulated the association between limbic network integration and memory consolidation.

Alterations of functional connectivity between the hippocampus and neocortex in aging may impact memory reactivations and the reorganization of memory traces within neocortical areas during sleep. Our results showed a positive association between memory consolidation and the functional centrality of the anterior hippocampus (importance of this region based on the strength of its connections), but not of the posterior hippocampus, supporting the specific role of this subregion in memory tasks involving objects (Cowan et al., 2020). The dichotomy within the hippocampus suggests that its anterior part is associated with representations of object-item and global contexts, whereas the posterior part is rather implicated in spatial information processing and navigation (Grady, 2020; Palacio & Cardenas, 2019; Poppenk et al., 2013). Performing an object-location memory task requires high associative learning abilities, supported by the anterior hippocampus, which is crucial for binding information (Grady, 2020; Jackson & Schacter, 2004; Poppenk et al., 2013). Age-related changes in the FC of the hippocampal subregions may explain the positive association between the centrality of the anterior hippocampus and memory consolidation. The anterior hippocampus exhibits stronger connectivity with frontal, temporal and medial orbitofrontal areas, whereas the posterior hippocampus is rather connected to the medial frontal and parietal cortices (Ranganath & Ritchey, 2012; Ritchey et al., 2015). Studies have shown a differential effect of age on the FC of hippocampal subregions, with most studies indicating a decrease in the posterior hippocampal FC, whereas the anterior hippocampal FC is less affected by age (Damoiseaux et al., 2016; Panitz et al., 2021). An increase in the FC of the posterior hippocampus in aging has also been observed, potentially resulting from a functional reorganization of anterior/posterior hippocampal connectivity with the neocortex and associated with a compensatory process (Blum et al., 2014).

Considering the functional brain reorganization that occurs with aging, we investigated the associations between memory consolidation and integration/segregation at both whole-brain and network levels. No associations were found at the whole-brain level, probably due to the fact that all networks are not equally affected by age, associative brain networks being particularly vulnerable (Betzel et al., 2014; Cassady et al., 2021; Chan et al., 2014; Damoiseaux, 2017; Geerligs et al., 2015; Malagurski et al., 2020; Spreng et al., 2016). We found that greater integration of the limbic network was associated with poorer overnight memory consolidation. This finding is consistent with the dedifferentiation hypothesis, which proposes that increased functional integration between networks can result from an age-related decrease in functional segregation (Koen & Rugg, 2019). Both functional segregation and integration follow a non-linear trajectory with aging, with inflection points occurring in the third and fourth decades, respectively (Deery et al., 2023). To determine whether the association between limbic network integration and memory consolidation was related to whole-brain segregation measures, analyses were controlled for modularity and system segregation. This association remained significant, suggesting that network integration *per se* predicts memory performance. Age-related functional reorganizations may lead to differences in network communication and contribute to lower cognitive performance, particularly for higher-order processes (Chan et al., 2014; Deery et al., 2023). Consistent with these studies, our results indicated that lower specialization, in particular a higher integration of the limbic network, is detrimental for memory consolidation, possibly contributing to the reduced memory performance with age. In the Schaefer atlas that was used in the present study, the limbic network includes orbitofrontal regions, the temporal pole and ventral anterior temporal lobe. These regions are connected by the uncinate fasciculus, which plays an important role in associative processes and episodic memory (Von Der Heide et al., 2013). The integrity of the uncinate fasciculus was found to mediate the effect of age on episodic memory (Merenstein et al., 2021). The disconnection of this tract was hypothesized to disrupt neural signals transmission contributing to memory impairment in older adults (Merenstein et al., 2021). The orbitofrontal cortex is known to be involved in memory formation (Ranganath et al., 2005), visuospatial memory (Cansino et al., 2015, 2017), as well as top-down cognitive and attentional control (Cansino et al., 2017). The anterior temporal lobe has been associated with perception of object properties and the binding between these properties (Ranganath & Ritchey, 2012; Ritchey et al., 2015). These regions subserve the cognitive processes involved in the object-location task used in this study.

Our analyses also revealed that, in participants with more fast spindles per train, the association between better memory consolidation and lower functional integration of the limbic network was stronger than in the other participants. However, no significant association was found with the centrality of the anterior hippocampus. Sleep spindles are known to facilitate neuronal plasticity (Cairney et al., 2018) and their clustering into trains constitute stable sleep periods for memory consolidation (Boutin & Doyon, 2020). Although no direct measure of FC during sleep was used in this study, our results suggest that longer fast spindle trains and a higher specialization of the limbic network during quiet wakefulness could have facilitated the reorganization of memory traces during sleep. Thus, sleep spindles could be a relevant marker of the association between rsFC of the limbic network and memory consolidation in aging.

To our knowledge, this study is the first investigating the associations between functional organization (at the whole-brain, network and hippocampal levels) and overnight memory consolidation in older adults. This study has several strengths. All participants underwent a habituation night to accustom to the sleep recording device. Analyses were controlled for multiple confounders such as age, sex, education and AHI. Participants were rigorously selected with standardized neuropsychological evaluations and strict exclusion criteria for neurological and psychiatric disorders. However, this study has also some limitations. First, we employed regressions to assess associations between sleep, rsFC and memory, it is therefore not possible to infer causal relationships. Second, although our statistical models included the AHI as a covariate, sleep apnea may affect memory consolidation and functional connectivity. Third, the absence of a wake group limits our ability to determine whether the observed effects are sleep-dependent or simply time-dependent. Finally, the absence of a young adult group prevents us from identifying age-related differences in rsFC and memory consolidation. Further studies should investigate the association between rsFC and memory consolidation using other sleep parameters, diverse memory tasks and participants whose age covers the whole adult lifespan.

## Conclusion

The present study provides evidence that the functional organization of the brain, particularly network integration and hippocampal centrality, plays a significant role in sleep-dependent memory consolidation in older adults. The higher specialization of the limbic network is beneficial to memory consolidation, supporting the dedifferentiation hypothesis of cognitive aging. The modulation of the relationship between the limbic network and overnight memory consolidation by fast spindle trains highlights the importance of sleep microstructure in cognitive aging. Future research should identify factors beyond age that influence the dynamics of sleep spindles, and determine how to promote hippocampal-cortical connectivity and optimize the functional specialization of brain networks. Such approaches may provide promising strategies to improve memory consolidation in older adults.

## Acknowledgements

The authors would like to thank Sebastien Polvent, Nicolas Oulhaj and Franck Doidy for their help with data acquisition. We acknowledge all the participants and the members of the Medit-Ageing Research Group (Alexandre Bejanin, Léa Chauveau, Anne Chocat, Fabienne Collette, Sophie Dautricourt, Robin De Flores, Marion Delarue, Hélène Espérou, Séverine Fauvel, Francesca Felisatti, Eglantine Ferrand Devouge, Eric Frison, Julie Gonneaud, Oriane Hébert, Olga Klimecki, Elizabeth Kuhn, Valérie Lefranc, Natalie Marchant, Cassandre Palix, Anne Quillard, Edelweiss Touron).

## Data availability statement

Data is available on request following a formal data sharing agreement and approval by the consortium and executive committee. The data sharing request form can be downloaded at https://silversantestudy.eu/2020/09/25/data-sharing/.

## Supplementary Material

**Table S1.** Resting-state functional connectivity characteristics.

| Graph measure | Mean ± SD [Min:Max] |
| --- | --- |
| **Whole-brain measures** |  |
| Global efficiency | 0.45 ± 0.08 [0.27:0.64] |
| Modularity (Q) | 0.35 ± 0.12 [0.12:0.53] |
| **Network measure: participation coefficient** |  |
| Dorsal attentional network | 0.68 ± 0.12 [0:0.81] |
| Salience/ventral network | 0.63 ± 0.19 [0:0.79] |
| Limbic network | 0.71 ± 0.21 [0:0.85] |
| Frontoparietal network | 0.58 ± 0.18 [0:0.77] |
| Default mode network | 0.48 ± 0.23 [0:0.73] |
| **Network measure: within-module degree z-score** |  |
| Dorsal attentional network | 0.08 ± 0.31 [-0.88:0.63] |
| Salience/ventral network | -0.04 ± 0.28 [-0.75:0.48] |
| Limbic network | 0.10 ± 0.37 [-0.73:0.56] |
| Frontoparietal network | -0.19 ± 0.27 [-0.59:0.46] |
| Default mode network | -0.17 ± 0.23 [-0.68:0.32] |
| **Hippocampal measure: hippocampal strength** |  |
| Anterior hippocampus | 15.90 ± 13.40 [0:56.5] |
| Posterior hippocampus | 11.10 ± 9.60 [0:38.9] |
| **Hippocampal measure: clustering coefficient** |  |
| Anterior hippocampus | 0.35 ± 0.18 [0:0.66] |
| Posterior hippocampus | 0.28 ± 0.19 [0:0.66] |

Data were obtained in 42 healthy older adults. Resting-state functional connectivity measures reported here were extracted at proportional threshold 15%. *SD*: Standard Deviation; *Max*: Maximum; *Min*: Minimum.

## Supplementary methods: graph analyses

Brain-network analyses were performed using functions implemented in the MATLAB-based Brain Connectivity Toolbox (Rubinov & Sporns, 2010). Whole-brain, network and ROI measures that were extracted are represented in **Figure 3C**.

## Whole-brain measures

Global efficiency and modularity (Q) measures were calculated to assess whole-brain integration and segregation, respectively.

Global efficiency is a measure of the integration of information between brain regions, quantifying the quality of information exchange across the brain. This measure was computed as the average length of the shortest inverse path between all nodes in the graph. An efficient system is characterized by short distances, with a small number of ROIs through which the information should pass to reach another ROI. A high global efficiency measure indicates a high degree of integrated and effective information transfer across the whole-brain (Rubinov & Sporns, 2010). The global efficiency metric of a graph *G* was defined as follows:

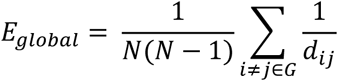

where N is the number of ROIs in the graph and *d_ij_* is the shortest path length between two ROIs *i* and *j*, N is the number of ROIs in the whole-brain.

The modularity statistic *Q* is a measure of segregation that indicates the balance between FC within- and between-networks across the whole-brain (Newman, 2006). Brain modularity (Q) was estimated based on the predefined networks using the Louvain algorithm (Blondel et al., 2008).

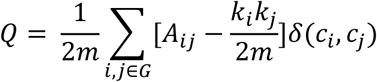

where *A_ij_* represents the weight of the connection between *i* and *j*, *k_i_* = ∑_*j*_ *A_ij_* is the sum of the weights of all connections attached to ROI *i*, *c_i_* is the predefined network to which ROI *i* is assigned, the *δ* function *δ(c_i_, c_j_)* equal 1 if *i* and *j* belong to the same predefined network and 0 otherwise and m = ½∑_*ij*_*A_ij_*. High segregation between networks is associated with high modularity (Q) value, while low modularity (Q) value indicates low segregation.

## Network measures

The participation coefficient and the within-module degree were calculated to characterize the network integration and segregation of the networks, respectively (Guimerà & Nunes Amaral, 2005; Rubinov & Sporns, 2010).

The participation coefficient (measure of integration or between-networks connectivity) quantifies the degree of connection of a ROI with ROIs assigned to other networks (Guimerà & Nunes Amaral, 2005). A high participation coefficient indicates that a ROI is strongly integrated across multiple distinct networks.

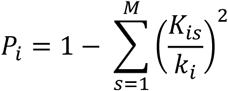

where *k_is_* is the connectivity weight between ROI *i* and other ROIs in network *s*, *k_i_* is the weight of all its connections and *M* is the number of networks in the whole-brain.

The within-module degree z-score (measure of segregation or within-network connectivity) indicates the strength of a ROI connectivity with the other ROIs in its network (Guimerà & Nunes Amaral, 2005).

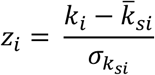

where *k_i_* is the connectivity weight between ROI *i* and other ROIs in network *s*, *k̅_si_* is the average of *k* over all of the ROIs in *s_i_*, and σ*_ksi_* is the standard deviation of *k* in *s_i_*, *z_i_* is called z-score.

To obtain a unique measure of integration and segregation per network, the median of the ROIs within each network was calculated for the measures of participation coefficient and within-module degree. As the average within-module degree is always equal to 0 for each network, the median allows an assessment of the degree of integration and segregation of networks

## Hippocampal measures

To estimate the centrality of the anterior and posterior hippocampus in the whole-brain, two measures of FC were calculated: hippocampal strength and clustering coefficient. The hippocampal strength measure quantifies the centrality of the hippocampus based on the strength of connectivity between the hippocampus and all other ROIs in the whole-brain. The hippocampal strength was computed as the sum of weighted connections of a ROI (Rubinov & Sporns, 2010).

The clustering coefficient is defined as a measure of segregation that represents the extent to which the interconnection of the hippocampus with its neighbor forms a cluster (Rubinov & Sporns, 2010).

The weighted clustering coefficient *C_i_* of the hippocampus *i* was calculated as:

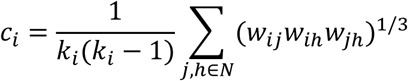

where *w_ij_*, *w_ih_*, and *w_jh_* are the connectivity weight between ROIs *i* and *j*, *i* and *h*, and *j* and *h*, respectively, *k_i_* is the number of connections of the hippocampus *i* and *N* is the set of all ROIs in the whole-brain.

## Supplementary methods and results: complementary analyses

Given graph measures were shown to provide complementary information about brain properties and that changes in one measure may represent a compensatory phenomenon or be a consequence of changes in another measure (Koen & Rugg, 2019), we conducted complementary analyses for all significant associations between memory consolidation and graph measures (anterior hippocampal ROI strength and participation coefficient of the limbic network). In these analyses, the other graph measure was added as covariate to consider potential interdependencies (**Table S2**).

**Table S2.** Summary of significant associations between memory consolidation and graph measures adjusted for another significant graph measure.

| Dependent variable | Independent variable | $\beta$ | SE | 95% CI | p |
| --- | --- | --- | --- | --- | --- |
| Overnight memory consolidation | Limbic network participation coefficient <sup>a</sup> | **-0.40** | **0.16** | **[-0.71:-0.08]** | **0.02** |
|  | Anterior hippocampal strength <sup>b</sup> | **0.33** | **0.16** | **[0.00:0.65]** | **0.049** |

Bold p-values were significant at $p < 0.05$ . <sup>a</sup> Model adjusted for age, sex, education, AHI, and anterior hippocampal strength. <sup>b</sup> Model adjusted for age, sex, education, AHI, and limbic network participation coefficient. Resting-state functional connectivity measures reported in this table were extracted at proportional threshold 15%. *CI: Confidence Interval; SE: Standard Error.*

Additionally, a regression analysis was performed to examine the association between the limbic network participation coefficient and anterior hippocampal strength (**Table S3**).

**Table S3.**
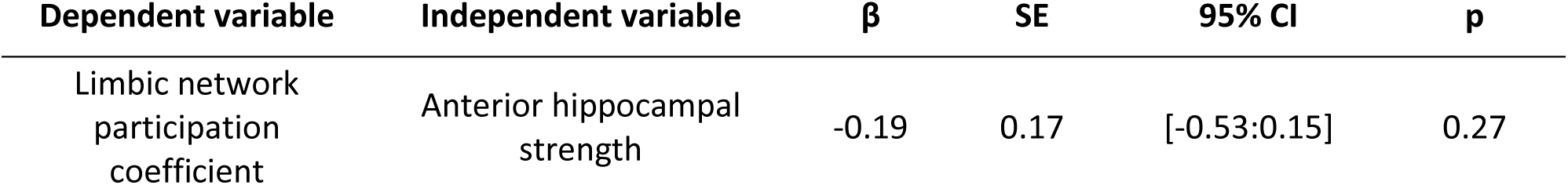
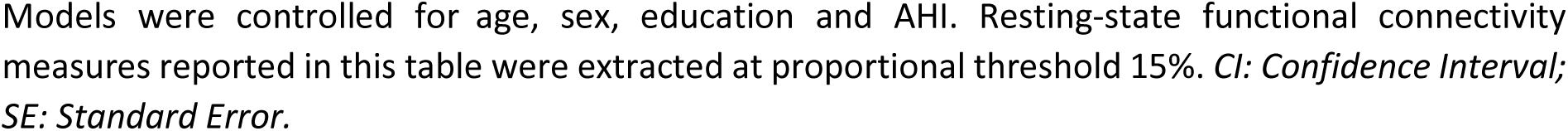
Association between the limbic network integration (participation coefficient) and functional centrality (strength) of the anterior hippocampus.

| Dependent variable | Independent variable | $\beta$ | SE | 95% CI | p |
| --- | --- | --- | --- | --- | --- |
| Limbic network participation coefficient | Anterior hippocampal strength | -0.19 | 0.17 | [-0.53:0.15] | 0.27 |

Models were controlled for age, sex, education and AHI. Resting-state functional connectivity measures reported in this table were extracted at proportional threshold 15%. *CI: Confidence Interval; SE: Standard Error.*

The lack of a significant association between the limbic network participation coefficient and anterior hippocampal strength (**Table S3**), combined with their independent relationships to memory consolidation (**Table S2**), suggests that these two graph measures may be related to sleep-dependent memory consolidation through distinct mechanisms.

**Table S4.** Summary of multiple linear regression models between overnight memory consolidation and graph measures (whole-brain, networks and hippocampus).

| Dependent variable | Independent variable | $\beta$ | SE | 95% CI | p |
| --- | --- | --- | --- | --- | --- |
| Overnight memory consolidation | **Whole-brain measures** |  |  |  |  |
|  | Global efficiency | -0.22 | 0.17 | [-0.58:0.13] | 0.21 |
|  | Modularity (Q) | -0.28 | 0.18 | [-0.65:0.10] | 0.14 |
|  | **Network measure: participation coefficient** |  |  |  |  |
|  | Dorsal attentional network | -0.26 | 0.17 | [-0.60:0.08] | 0.13 |
|  | Salience/ventral network | -0.04 | 0.18 | [-0.40:0.32] | 0.81 |
|  | Limbic network | **-0.46** | **0.16** | **[-0.78:-0.13]** | **0.007\*** |
|  | Frontoparietal network | -0.15 | 0.17 | [-0.50:0.19] | 0.38 |
|  | Default mode network | **-0.39** | **0.17** | **[-0.74:-0.04]** | **0.03** |
|  | Visual network | 0.03 | 0.17 | [-0.74:-0.04] | 0.86 |
|  | Sensorimotor network | 0.16 | 0.17 | [0.17:-0.19] | 0.36 |
|  | **Network measure: within-module degree** |  |  |  |  |
|  | Dorsal attentional network | -0.23 | 0.17 | [-0.57:0.11] | 0.18 |
|  | Salience/ventral network | -0.05 | 0.17 | [-0.40:0.30] | 0.77 |
|  | Limbic network | **-0.42** | **0.16** | **[-0.75:-0.10]** | **0.01** |
|  | Frontoparietal network | 0.04 | 0.17 | [-0.30:0.39] | 0.80 |
|  | Default mode network | -0.02 | 0.19 | [-0.40:0.36] | 0.91 |
|  | Visual network | 0.08 | 0.17 | [-0.27:0.43] | 0.66 |
|  | Sensorimotor network | -0.06 | 0.17 | [-0.41:0.29] | 0.73 |
|  | **Hippocampal measure: hippocampal strength** |  |  |  |  |
|  | Anterior hippocampus | **0.40** | **0.17** | **[0.06:0.74]** | **0.02\*** |
|  | Posterior hippocampus | 0.02 | 0.17 | [-0.33:0.37] | 0.90 |
|  | **Hippocampal measure: clustering coefficient** |  |  |  |  |
|  | Anterior hippocampus | 0.07 | 0.18 | [-0.30:0.44] | 0.70 |
|  | Posterior hippocampus | -0.04 | 0.17 | [-0.38:0.31] | 0.84 |

Bold p-values were significant at $p < 0.05$ and \* survived FDR correction. Statistical models were controlled by age, sex, education and AHI. Resting-state functional connectivity measures reported in this table were extracted at proportional threshold 15%. *CI: Confidence Interval; SE: Standard Error.*

**Table S5.** Summary of multiple linear regression models of limbic network integration and memory consolidation adjusted for whole-brain segregation.

| Dependent variable | Independent variable | $\beta$ | SE | 95% CI | p |
| --- | --- | --- | --- | --- | --- |
| Overnight memory consolidation | Network measure: participation coefficient |  |  |  |  |
|  | Limbic network <sup>a</sup> | **-0.44** | **0.16** | **[-0.76:-0.12]** | **0.009** |
|  | Limbic network <sup>b</sup> | **-0.43** | **0.17** | **[-0.77:-0.09]** | **0.01** |

Bold p-values were significant at $p < 0.05$ . Statistical models were controlled by <sup>a</sup> age, sex, education, AHI and whole brain modularity (Q) or by <sup>b</sup> age, sex, education, AHI and system segregation (Chan et al., 2014). Resting-state functional connectivity measures reported in this table were extracted at proportional threshold 15%. *CI: Confidence Interval; SE: Standard Error.*

**Table S6.** Summary of multiple linear regression models of the modulation effect of fast spindle train on associations between resting-state functional connectivity and memory consolidation.

| Dependent variable | Independent variable | $\beta$ | SE | 95% CI | p |
| --- | --- | --- | --- | --- | --- |
| Limbic network participation coefficient | Overnight memory consolidation * mean number of fast spindle per train | **-3.45** | **0.03** | **[-3.46:-3.44]** | **0.02** |
| Anterior hippocampal strength |  | -0.73 | 2.01 | [-0.73:0.45] | 0.63 |

Bold p-values were significant at $p < 0.05$ . Models were controlled by age, sex, education, AHI. Resting-state functional connectivity measures reported in this table were extracted at proportional threshold 15%. *CI: Confidence Interval; SE: Standard Error.*

## References

1. Baltes, P. B., & Lindenberger, U. (1997). Emergence of a powerful connection between sensory and cognitive functions across the adult life span: A new window to the study of cognitive aging? Psychology and Aging, 12(1), 12–21. 10.1037/0882-7974.12.1.12

2. Benjamini, Y., & Hochberg, Y. (1995). Controlling the False Discovery Rate: A Practical and Powerful Approach to Multiple Testing. Journal of the Royal Statistical Society. Series B (Methodological), 57(1), 289–300.

3. Betzel, R. F., Byrge, L., He, Y., Goñi, J., Zuo, X.-N., & Sporns, O. (2014). Changes in structural and functional connectivity among resting-state networks across the human lifespan. NeuroImage, 102, 345–357. 10.1016/j.neuroimage.2014.07.067

4. Blum, S., Habeck, C., Steffener, J., Razlighi, Q., & Stern, Y. (2014). Functional connectivity of the posterior hippocampus is more dominant as we age. Cognitive Neuroscience, 5(3–4), 150–159. 10.1080/17588928.2014.975680

5. Born, J., & Wilhelm, I. (2012). System consolidation of memory during sleep. Psychological Research, 76(2), 192–203. 10.1007/s00426-011-0335-6

6. Boutin, A., & Doyon, J. (2020). A sleep spindle framework for motor memory consolidation. Philosophical Transactions of the Royal Society B: Biological Sciences, 375(1799), 20190232. 10.1098/rstb.2019.0232

7. Bullmore, E., & Sporns, O. (2009). Complex brain networks: Graph theoretical analysis of structural and functional systems. Nature Reviews Neuroscience, 10(3), 186–198. 10.1038/nrn2575

8. Buzsáki, G. (1996). The Hippocampo-Neocortical Dialogue. Cerebral Cortex, 6(2), 81–92. 10.1093/cercor/6.2.81

9. Cairney, S. A., Guttesen, A. Á. V., El Marj, N., & Staresina, B. P. (2018). Memory Consolidation Is Linked to Spindle-Mediated Information Processing during Sleep. Current Biology, 28(6), 948–954.e4. 10.1016/j.cub.2018.01.087

10. Cansino, S., Estrada-Manilla, C., Trejo-Morales, P., Pasaye-Alcaraz, E. H., Aguilar-Castañeda, E., Salgado-Lujambio, P., & Sosa-Ortiz, A. L. (2015). fMRI subsequent source memory effects in young, middle-aged and old adults. Behavioural Brain Research, 280, 24–35. 10.1016/j.bbr.2014.11.042

11. Cansino, S., Trejo-Morales, P., Estrada-Manilla, C., Pasaye-Alcaraz, E. H., Aguilar-Castañeda, E., Salgado-Lujambio, P., & Sosa-Ortiz, A. L. (2017). Effective connectivity during successful and unsuccessful recollection in young and old adults. Neuropsychologia, 103, 168–182. 10.1016/j.neuropsychologia.2017.07.016

12. Cao, M., Wang, J.-H., Dai, Z.-J., Cao, X.-Y., Jiang, L.-L., Fan, F.-M., Song, X.-W., Xia, M.-R., Shu, N., Dong, Q., Milham, M. P., Castellanos, F. X., Zuo, X.-N., & He, Y. (2014). Topological organization of the human brain functional connectome across the lifespan. Developmental Cognitive Neuroscience, 7, 76–93. 10.1016/j.dcn.2013.11.004

13. Cassady, K. E., Adams, J. N., Chen, X., Maass, A., Harrison, T. M., Landau, S., Baker, S., & Jagust, W. (2021). Alzheimer’s Pathology Is Associated with Dedifferentiation of Intrinsic Functional Memory Networks in Aging. Cerebral Cortex, 31(10), 4781–4793. 10.1093/cercor/bhab122

14. Champetier, P., André, C., Weber, F. D., Rehel, S., Ourry, V., Laniepce, A., Lutz, A., Bertran, F., Cabé, N., Pitel, A.-L., Poisnel, G., De La Sayette, V., Vivien, D., Chételat, G., & Rauchs, G. (2023). Age-related changes in fast spindle clustering during non-rapid eye movement sleep and their relevance for memory consolidation. SLEEP, 46(5), zsac282. 10.1093/sleep/zsac282

15. Chan, M. Y., Park, D. C., Savalia, N. K., Petersen, S. E., & Wig, G. S. (2014). Decreased segregation of brain systems across the healthy adult lifespan. Proceedings of the National Academy of Sciences, 111(46). 10.1073/pnas.1415122111

16. Cohen, J. R., & D’Esposito, M. (2016). The Segregation and Integration of Distinct Brain Networks and Their Relationship to Cognition. The Journal of Neuroscience, 36(48), 12083–12094. 10.1523/JNEUROSCI.2965-15.2016

17. Cowan, E., Liu, A., Henin, S., Kothare, S., Devinsky, O., & Davachi, L. (2020). Sleep Spindles Promote the Restructuring of Memory Representations in Ventromedial Prefrontal Cortex through Enhanced Hippocampal–Cortical Functional Connectivity. The Journal of Neuroscience, 40(9), 1909–1919. 10.1523/JNEUROSCI.1946-19.2020

18. Crowell, C. A., Davis, S. W., Beynel, L., Deng, L., Lakhlani, D., Hilbig, S. A., Palmer, H., Brito, A., Peterchev, A. V., Luber, B., Lisanby, S. H., Appelbaum, L. G., & Cabeza, R. (2020). Older adults benefit from more widespread brain network integration during working memory. NeuroImage, 218, 116959. 10.1016/j.neuroimage.2020.116959

19. Crowley, K. (2002). The effects of normal aging on sleep spindle and K-complex production. Clinical Neurophysiology, 113(10), 1615–1622. 10.1016/S1388-2457(02)00237-7

20. Damoiseaux, J. S. (2017). Effects of aging on functional and structural brain connectivity. NeuroImage, 160, 32–40. 10.1016/j.neuroimage.2017.01.077

21. Damoiseaux, J. S., Viviano, R. P., Yuan, P., & Raz, N. (2016). Differential effect of age on posterior and anterior hippocampal functional connectivity. NeuroImage, 133, 468–476. 10.1016/j.neuroimage.2016.03.047

22. Deery, H. A., Di Paolo, R., Moran, C., Egan, G. F., & Jamadar, S. D. (2023). The older adult brain is less modular, more integrated, and less efficient at rest: A systematic review of large-scale resting-state functional brain networks in aging. Psychophysiology, 60(1), e14159. 10.1111/psyp.14159

23. Deleglise, A., Donnelly-Kehoe, P. A., Yeffal, A., Jacobacci, F., Jovicich, J., Amaro Jr, E., Armony, J. L., Doyon, J., & Della-Maggiore, V. (2023). Human motor sequence learning drives transient changes in network topology and hippocampal connectivity early during memory consolidation. Cerebral Cortex, 33(10), 6120–6131. 10.1093/cercor/bhac489

24. Diekelmann, S., & Born, J. (2010). The memory function of sleep. Nature Reviews Neuroscience, 11(2), 114–126. 10.1038/nrn2762

25. Esposito, R., Cieri, F., Chiacchiaretta, P., Cera, N., Lauriola, M., Di Giannantonio, M., Tartaro, A., & Ferretti, A. (2018). Modifications in resting state functional anticorrelation between default mode network and dorsal attention network: Comparison among young adults, healthy elders and mild cognitive impairment patients. Brain Imaging and Behavior, 12(1), 127–141. 10.1007/s11682-017-9686-y

26. Fernandez, L. M. J., & Lüthi, A. (2020). Sleep Spindles: Mechanisms and Functions. Physiological Reviews, 100(2), 805–868. 10.1152/physrev.00042.2018

27. Ferreira, L. K., Regina, A. C. B., Kovacevic, N., Martin, M. D. G. M., Santos, P. P., Carneiro, C. D. G., Kerr, D. S., Amaro, E., McIntosh, A. R., & Busatto, G. F. (2016). Aging Effects on Whole-Brain Functional Connectivity in Adults Free of Cognitive and Psychiatric Disorders. Cerebral Cortex, 26(9), 3851–3865. 10.1093/cercor/bhv190

28. Fjell, A. M., Sneve, M. H., Grydeland, H., Storsve, A. B., De Lange, A.-M. G., Amlien, I. K., Røgeberg, O. J., & Walhovd, K. B. (2015). Functional connectivity change across multiple cortical networks relates to episodic memory changes in aging. Neurobiology of Aging, 36(12), 3255–3268. 10.1016/j.neurobiolaging.2015.08.020

29. Fjell, A. M., Sneve, M. H., Storsve, A. B., Grydeland, H., Yendiki, A., & Walhovd, K. B. (2016). Brain Events Underlying Episodic Memory Changes in Aging: A Longitudinal Investigation of Structural and Functional Connectivity. Cerebral Cortex, 26(3), 1272–1286. 10.1093/cercor/bhv102

30. Friedman, D., Nessler, D., & Johnson, R. (2007). Memory Encoding and Retrieval in the Aging Brain. Clinical EEG and Neuroscience, 38(1), 2–7. 10.1177/155005940703800105

31. Gallen, C. L., & D’Esposito, M. (2019). Brain Modularity: A Biomarker of Intervention-related Plasticity. Trends in Cognitive Sciences, 23(4), 293–304. 10.1016/j.tics.2019.01.014

32. Geerligs, L., Renken, R. J., Saliasi, E., Maurits, N. M., & Lorist, M. M. (2015). A Brain-Wide Study of Age-Related Changes in Functional Connectivity. Cerebral Cortex, 25(7), 1987–1999. 10.1093/cercor/bhu012

33. Grady, C. (2020). Meta-analytic and functional connectivity evidence from functional magnetic resonance imaging for an anterior to posterior gradient of function along the hippocampal axis. Hippocampus, 30(5), 456–471. 10.1002/hipo.23164

34. Grady, C., Sarraf, S., Saverino, C., & Campbell, K. (2016). Age differences in the functional interactions among the default, frontoparietal control, and dorsal attention networks. Neurobiology of Aging, 41, 159–172. 10.1016/j.neurobiolaging.2016.02.020

35. Hamel, A., Mary, A., & Rauchs, G. (2023). Sleep and memory consolidation in aging: A neuroimaging perspective. Revue Neurologique, 179(7), 658–666. 10.1016/j.neurol.2023.08.003

36. Huo, L., Li, R., Wang, P., Zheng, Z., & Li, J. (2018). The Default Mode Network Supports Episodic Memory in Cognitively Unimpaired Elderly Individuals: Different Contributions to Immediate Recall and Delayed Recall. Frontiers in Aging Neuroscience, 10, 6. 10.3389/fnagi.2018.00006

37. Jackson, O., & Schacter, D. L. (2004). Encoding activity in anterior medial temporal lobe supports subsequent associative recognition. NeuroImage, 21(1), 456–462. 10.1016/j.neuroimage.2003.09.050

38. Jacobs, H. I. L., Dillen, K. N. H., Risius, O., GÃ¶reci, Y., Onur, O. A., Fink, G. R., & Kukolja, J. (2015). Consolidation in older adults depends upon competition between resting-state networks. Frontiers in Aging Neuroscience, 6. 10.3389/fnagi.2014.00344

39. Klinzing, J. G., Niethard, N., & Born, J. (2019). Mechanisms of systems memory consolidation during sleep. Nature Neuroscience, 22(10), 1598–1610. 10.1038/s41593-019-0467-3

40. Koen, J. D., & Rugg, M. D. (2019). Neural Dedifferentiation in the Aging Brain. Trends in Cognitive Sciences, 23(7), 547–559. 10.1016/j.tics.2019.04.012

41. Kukolja, J., Göreci, D. Y., Onur, Ö. A., Riedl, V., & Fink, G. R. (2016). Resting-state fMRI evidence for early episodic memory consolidation: Effects of age. Neurobiology of Aging, 45, 197–211. 10.1016/j.neurobiolaging.2016.06.004

42. Landolt, H.-P., Dijk, D.-J., Achermann, P., & Borbély, A. A. (1996). Effect of age on the sleep EEG: Slow-wave activity and spindle frequency activity in young and middle-aged men. Brain Research, 738(2), 205–212. 10.1016/S0006-8993(96)00770-6

43. Luppi, A. I., Craig, M. M., Pappas, I., Finoia, P., Williams, G. B., Allanson, J., Pickard, J. D., Owen, A. M., Naci, L., Menon, D. K., & Stamatakis, E. A. (2019). Consciousness-specific dynamic interactions of brain integration and functional diversity. Nature Communications, 10(1), Article 1. 10.1038/s41467-019-12658-9

44. Malagurski, B., Liem, F., Oschwald, J., Mérillat, S., & Jäncke, L. (2020). Functional dedifferentiation of associative resting state networks in older adults – A longitudinal study. NeuroImage, 214, 116680. 10.1016/j.neuroimage.2020.116680

45. Merenstein, J. L., Corrada, M. M., Kawas, C. H., & Bennett, I. J. (2021). Age affects white matter microstructure and episodic memory across the older adult lifespan. Neurobiology of Aging, 106, 282–291. 10.1016/j.neurobiolaging.2021.06.021

46. Muehlroth, B. E., Rasch, B., & Werkle-Bergner, M. (2020). Episodic memory consolidation during sleep in healthy aging. Sleep Medicine Reviews, 52, 101304. 10.1016/j.smrv.2020.101304

47. Muehlroth, B. E., Sander, M. C., Fandakova, Y., Grandy, T. H., Rasch, B., Lee Shing, Y., & Werkle-Bergner, M. (2020). Memory quality modulates the effect of aging on memory consolidation during sleep: Reduced maintenance but intact gain. NeuroImage, 209, 116490. 10.1016/j.neuroimage.2019.116490

48. Nicolas, A., Petit, D., Rompré, S., & Montplaisir, J. (2001). Sleep spindle characteristics in healthy subjects of different age groups. Clinical Neurophysiology, 112(3), 521–527. 10.1016/S1388-2457(00)00556-3

49. Nyberg, L. (2017). Functional brain imaging of episodic memory decline in ageing. Journal of Internal Medicine, 281(1), 65–74. 10.1111/joim.12533

50. Onoda, K., & Yamaguchi, S. (2013). Small-worldness and modularity of the resting-state functional brain network decrease with aging. Neuroscience Letters, 556, 104–108. 10.1016/j.neulet.2013.10.023

51. Palacio, N., & Cardenas, F. (2019). A systematic review of brain functional connectivity patterns involved in episodic and semantic memory. Reviews in the Neurosciences, 30(8), 889–902. 10.1515/revneuro-2018-0117

52. Panitz, D. Y., Berkovich-Ohana, A., & Mendelsohn, A. (2021). Age-related functional connectivity along the hippocampal longitudinal axis. Hippocampus, 31(10), 1115–1127. 10.1002/hipo.23377

53. Pedersen, R., Geerligs, L., Andersson, M., Gorbach, T., Avelar-Pereira, B., Wåhlin, A., Rieckmann, A., Nyberg, L., & Salami, A. (2021). When functional blurring becomes deleterious: Reduced system segregation is associated with less white matter integrity and cognitive decline in aging. NeuroImage, 242, 118449. 10.1016/j.neuroimage.2021.118449

54. Poisnel, G., Arenaza-Urquijo, E., Collette, F., Klimecki, O. M., Marchant, N. L., Wirth, M., De La Sayette, V., Rauchs, G., Salmon, E., Vuilleumier, P., Frison, E., Maillard, A., Vivien, D., Lutz, A., Chételat, G., & Medit-Ageing Research Group. (2018). The Age-Well randomized controlled trial of the Medit-Ageing European project: Effect of meditation or foreign language training on brain and mental health in older adults. Alzheimer’s & Dementia: Translational Research & Clinical Interventions, 4(1), 714–723. 10.1016/j.trci.2018.10.011

55. Poppenk, J., Evensmoen, H. R., Moscovitch, M., & Nadel, L. (2013). Long-axis specialization of the human hippocampus. Trends in Cognitive Sciences, 17(5), 230–240. 10.1016/j.tics.2013.03.005

56. Ranganath, C., Heller, A., Cohen, M. X., Brozinsky, C. J., & Rissman, J. (2005). Functional connectivity with the hippocampus during successful memory formation. Hippocampus, 15(8), 997–1005. 10.1002/hipo.20141

57. Ranganath, C., & Ritchey, M. (2012). Two cortical systems for memory-guided behaviour. Nature Reviews Neuroscience, 13(10), 713–726. 10.1038/nrn3338

58. Rasch, B., Büchel, C., Gais, S., & Born, J. (2007). Odor Cues During Slow-Wave Sleep Prompt Declarative Memory Consolidation. Science, 315(5817), 1426–1429. 10.1126/science.1138581

59. Reuter-Lorenz, P. A., & Lustig, C. (2005). Brain aging: Reorganizing discoveries about the aging mind. Current Opinion in Neurobiology, 15(2), 245–251. 10.1016/j.conb.2005.03.016

60. Ritchey, M., Libby, L. A., & Ranganath, C. (2015). Cortico-hippocampal systems involved in memory and cognition. In Progress in Brain Research (Vol. 219, pp. 45–64). Elsevier. 10.1016/bs.pbr.2015.04.001

61. Rubinov, M., & Sporns, O. (2010). Complex network measures of brain connectivity: Uses and interpretations. NeuroImage, 52(3), 1059–1069. 10.1016/j.neuroimage.2009.10.003

62. Sala-Llonch, R. (2014). Changes in whole-brain functional networks and memory performance in aging. Neurobiology of Aging.

63. Schaefer, A., Kong, R., Gordon, E. M., Laumann, T. O., Zuo, X.-N., Holmes, A. J., Eickhoff, S. B., & Yeo, B. T. T. (2018). Local-Global Parcellation of the Human Cerebral Cortex from Intrinsic Functional Connectivity MRI. *Cerebral Cortex (New York*, NY), 28(9), 3095–3114. 10.1093/cercor/bhx179

64. Shaw, E. E., Schultz, A. P., Sperling, R. A., & Hedden, T. (2015). Functional Connectivity in Multiple Cortical Networks Is Associated with Performance Across Cognitive Domains in Older Adults. Brain Connectivity, 5(8), 505–516. 10.1089/brain.2014.0327

65. Song, J., Birn, R. M., Meyerand, M. E., & Prabhakaran, V. (2014). Age-Related Reorganizational Changes in Modularity and Functional Connectivity of Human Brain Networks. Brain Connect., 4(9), 662–676. 10.1089/brain.2014.0286

66. Sonni, A., & Spencer, R. M. C. (2015). Sleep protects memories from interference in older adults. Neurobiology of Aging, 36(7), 2272–2281. 10.1016/j.neurobiolaging.2015.03.010

67. Sporns, O., & Betzel, R. F. (2016). Modular Brain Networks. Annual Review of Psychology, 67(1), 613– 640. 10.1146/annurev-psych-122414-033634

68. Spreng, R. N., Stevens, W. D., Viviano, J. D., & Schacter, D. L. (2016). Attenuated anticorrelation between the default and dorsal attention networks with aging: Evidence from task and rest. Neurobiology of Aging, 45, 149–160. 10.1016/j.neurobiolaging.2016.05.020

69. Stumme, J., Jockwitz, C., Hoffstaedter, F., Amunts, K., & Caspers, S. (2020). Functional network reorganization in older adults: Graph-theoretical analyses of age, cognition and sex. NeuroImage, 214, 116756. 10.1016/j.neuroimage.2020.116756

70. Tian, Y., Margulies, D. S., Breakspear, M., & Zalesky, A. (2020). Topographic organization of the human subcortex unveiled with functional connectivity gradients. Nature Neuroscience, 23(11), 1421– 1432. 10.1038/s41593-020-00711-6

71. Tromp, D., Dufour, A., Lithfous, S., Pebayle, T., & Després, O. (2015). Episodic memory in normal aging and Alzheimer disease: Insights from imaging and behavioral studies. Ageing Research Reviews, 24, 232–262. 10.1016/j.arr.2015.08.006

72. Villain, N., Landeau, B., Groussard, M., Mevel, K., Fouquet, M., Dayan, J., Eustache, F., Desgranges, B., & Chételat, G. (2010). A Simple Way to Improve Anatomical Mapping of Functional Brain Imaging. Journal of Neuroimaging, 20(4), 324–333. 10.1111/j.1552-6569.2010.00470.x

73. Von Der Heide, R. J., Skipper, L. M., Klobusicky, E., & Olson, I. R. (2013). Dissecting the uncinate fasciculus: Disorders, controversies and a hypothesis. Brain, 136(6), 1692–1707. 10.1093/brain/awt094

74. Wig, G. S. (2017). Segregated Systems of Human Brain Networks. Trends in Cognitive Sciences, 21(12), 981–996. 10.1016/j.tics.2017.09.006

75. Wilson, J. K., Baran, B., Pace-Schott, E. F., Ivry, R. B., & Spencer, R. M. C. (2012). Sleep modulates word-pair learning but not motor sequence learning in healthy older adults. Neurobiology of Aging, 33(5), 991–1000. 10.1016/j.neurobiolaging.2011.06.029

76. Wu, J., Li, J., Eickhoff, S. B., Hoffstaedter, F., Hanke, M., Yeo, B. T. T., & Genon, S. (2022). Cross-cohort replicability and generalizability of connectivity-based psychometric prediction patterns. NeuroImage, 262, 119569. 10.1016/j.neuroimage.2022.119569

77. Xu, K., Liu, Y., Zhan, Y., Ren, J., & Jiang, T. (2018). BRANT: A Versatile and Extendable Resting-State fMRI Toolkit. Frontiers in Neuroinformatics, 12, 52. 10.3389/fninf.2018.00052

78. Zalesky, A., Fornito, A., Cocchi, L., Gollo, L. L., Van Den Heuvel, M. P., & Breakspear, M. (2016). Connectome sensitivity or specificity: Which is more important? NeuroImage, 142, 407–420. 10.1016/j.neuroimage.2016.06.035

## References

80. Blondel, V. D., Guillaume, J.-L., Lambiotte, R., & Lefebvre, E. (2008). Fast unfolding of communities in large networks. Journal of Statistical Mechanics: Theory and Experiment, 2008(10), P10008. 10.1088/1742-5468/2008/10/P10008

81. Guimerà, R., & Nunes Amaral, L. A. (2005). Functional cartography of complex metabolic networks. Nature, 433(7028), 895–900. 10.1038/nature03288

82. Newman, M. E. J. (2006). Finding community structure in networks using the eigenvectors of matrices. Physical Review E, 74(3), 036104. 10.1103/PhysRevE.74.036104

83. Rubinov, M., & Sporns, O. (2010). Complex network measures of brain connectivity: Uses and interpretations. NeuroImage, 52(3), 1059–1069. 10.1016/j.neuroimage.2009.10.003

